# Stimulus predictability does not modulate bottom-up attentional capture

**DOI:** 10.1101/292508

**Authors:** Erik L. Meijs, Felix H. Klaassen, Levan Bokeria, Simon van Gaal, Floris P. de Lange

## Abstract

Attention can be involuntarily captured by physically salient stimuli, a phenomenon known as bottom-up attention. Typically, these salient stimuli occur unpredictably in time and space. Therefore, in a series of three behavioral experiments, we investigated the extent to which such bottom-up attentional capture is a function of one’s prior expectations. In the context of an exogenous cueing task, we systematically manipulated participants’ spatial (Experiment 1) or temporal (Experiment 2 and 3) expectations about an uninformative cue, and examined the extent of attentional capture by the cue. We anticipated larger attentional capture for unexpected compared to expected cues. However, while we observed robust attentional capture in all experiments, we did not find any evidence for a modulation of attentional capture by prior expectation. This underscores the automatic and reflexive nature of bottom-up attention.

## Introduction

When performing tasks in our everyday lives, we constantly have to battle potential distraction by task-irrelevant inputs. Even when we want to stay focused on the task at hand, it can be difficult to ignore other, often more salient, stimuli that capture our attention. Historically, there has been considerable debate on whether attentional capture is purely stimulus-driven [1,2], or also depends on top-down goals [3]. More recently it has been suggested that recent trial history [4] and associations with reward [5,6] may also strongly modulate attentional capture. This implies that the presence and amount of attentional capture may be a complex function of both stimulus and internal variables [7].

Stimulus expectation is another factor that may modulate bottom-up capture. One of the studies providing evidence for this was carried out by Folk and Remington [8]. In a spatial cueing paradigm, they manipulated the frequency of salient but uninformative (i.e., not predictive of the target) cues. Their results indicated that these cues captured attention only when they were unlikely, regardless of the top-down task set participants were using (but see [9] for an alternative interpretation). In another study the proportion of distractors was systematically varied over blocks [10]. The distractors interfered more with target processing when they were presented in a block with fewer distractors, suggesting they captured attention more when they were more surprising. Similarly, it has been observed that novel stimuli are most potent in capturing attention [11] and also most robustly modulate the neural response in a macaque’s V1 [12]. Taken together, these studies support the hypothesis that attentional capture by task-irrelevant stimuli may be modulated by perceptual expectations, and most notably by the violation of these expectations. This can be interpreted as evidence that surprising stimuli are more salient and therefore more attention-grabbing [13,14]. Additional evidence supporting this idea comes from studies on mismatch detection, in which it has been shown that unexpected deviant stimuli lead to larger mismatch responses in the EEG-signal and seem to subjectively “pop out” [15–17].

Besides influencing the amount of attentional capture by distracting stimuli, there is evidence that prior information about these cues can help participants to voluntarily diminish distraction [18,19]. For example, it has been shown that attentional capture by unlikely distractors can be attenuated when the search task promotes suppressing features similar to those of the distractors stimuli [9]. Another study showed that it is easier to ignore regular sequences than irregular sequences [20]. Whether reducing the amount of distraction is caused by the inhibition of attentional capture, or by rapid disengagement at a later stage is still debated [4,21]. An electrophysiological study by Kiss and colleagues suggested that bottom-up capture can be inhibited, but that this only happens when task demands (i.e., timing) require it [22].

One interpretation of the empirical evidence above is that surprising stimuli are more salient and therefore more attention-grabbing [13,14]. Predictive coding theories have suggested that processing unexpected events requires more resources [23,24]. One may conceptualize bottom-up attention as a way of redistributing resources, for example towards processing unexpected events. This is in line with findings that bottom-up attention increases contrast sensitivity [25]. Nevertheless, many models based on predictive coding have actually suggested that regularity and predictability may attract attention [26]. The idea behind this is that predictable inputs are more strongly weighted because they are more reliable. A number of studies have provided support for this idea [27], suggesting that regularities automatically attract attention (but see e.g. [28]).

It thus seems that the link between expectation and bottom-up attention is still far from clear [24,29]. Attention may be either drawn to surprising stimuli, or to regularly occurring ones. Therefore, we performed a series of three experiments using an exogenous cueing task [30], in which we explicitly looked at this relationship by manipulating participants’ expectations about an otherwise uninformative (i.e., unrelated to target) cue stimulus. Specifically, we investigated to which extent prior expectations about the cue modulated bottom-up attentional capture. Based on the evidence listed earlier, we anticipated that unexpected cue stimuli attract more attention and therefore in larger cue-target validity effects (i.e., performance difference between validly and invalidly cued trials), whereas expected cue stimuli are followed by strongly reduced or even absent validity effects. To preview, in contrast to this hypothesis, we observed attentional capture in all experiments, but no modulation by prior knowledge about the cue stimulus in any experiment.

## Methods

### Participants

We tested a total of 248 participants in a series of three experiments (120 participants in Experiment 1, 67 participants in Experiment 2 and 61 participants in Experiment 3). All participants had normal or corrected-to-normal vision.

We excluded 4 participants from Experiment 2 because button presses were not recorded properly. Furthermore, from each experiment we excluded participants who failed to respond on more than 20% of trials or whose task performance was markedly (more than 3 standard deviations) worse than that of other subjects (Experiment 1: 2 participants excluded; Experiment 3: 3 participants excluded). In the end, we included 118 participants for Experiment 1 (87 females, age 22.7±5.0 years), 63 participants for Experiment 2 (50 females, age 23.0±3.5 years) and 58 participants for Experiment 3 (42 females, age 22.7±3.5 years).

All experiments were approved by the local ethics committee of the Radboud University (CMO Arnhem-Nijmegen; 2014/288 “Imaging Human Cognition”). Written informed consent was obtained from all participants according to the Declaration of Helsinki. Compensation for each of the studies was either 8 Euros or course credit.

### Materials

All stimuli were presented using the Psychophysics Toolbox (Brainard, 1997) within MATLAB (MathWorks, Natick, MA, USA). Stimuli were generated by a Dell T3500 Workstation and displayed on a 24” BENQ LED monitor (1920 × 1080 pixels; 60Hz; screen size 53.1cm x 29.9 cm). All presented stimuli were “black” (RGB: [0 0 0]; ± 0.3 cd/m^2^) on a grey (RGB: [150 150 150]; ± 103.9 cd/m^2^) background. A chinrest was used in all experiments to control the distance participants were seated from the monitor (±57 cm). In Experiment 1 participants responded by means of two button boxes. For Experiments 2 and 3 participants used the computer keyboard (DELL KB522).

### Procedure and stimuli

In all experiments, participants performed an adjusted version of the exogenous cueing task (Posner, 1980; Figure 1A). First, a cue (2° circular outline) was presented for 50 ms either 5 degrees above or below fixation. After the cue, a target was presented centered on either same (valid trials) or the opposite (invalid trials) screen location. Cue location and target location were unrelated, meaning that both target locations were equally likely throughout the experiment, regardless of where the cue was presented. Targets were small (0.48° wide and 0.60° high) arrows pointing either leftward or rightward. The participants’ task was to report the direction the arrow was pointing in (leftward or rightward) by pressing a button with either their left or right index finger, while maintaining fixation throughout the experiment.

**Figure 1.**
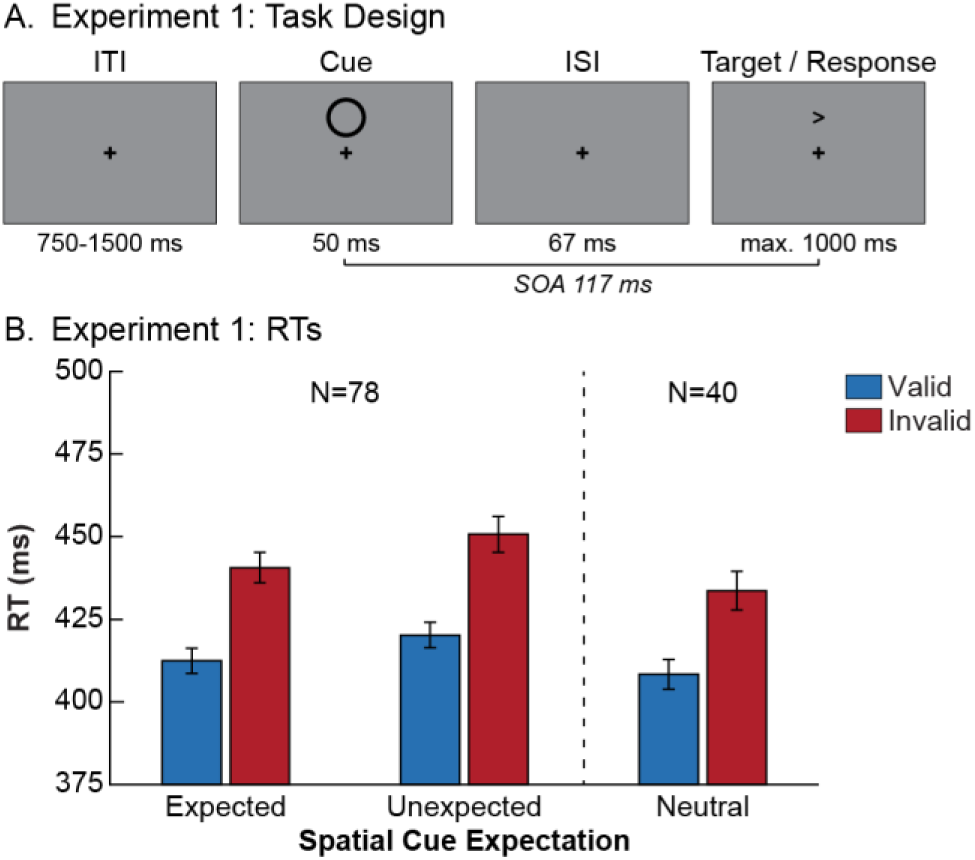
Task design and behavioral results of Experiment 1. **(A)** Trial structure of the exogenous cueing task used in Experiment 1. In every trial a cue (circular annulus) was presented for 50 ms, either above or below fixation. After an inter stimulus interval (ISI) of 67 ms (SOA 117 ms), a target (arrow) was presented in either the same (valid trials) or opposite (invalid trials) location. We manipulated spatial cue expectation by varying the likelihood the cue would appear in either location. In one group of participants the cue appeared equally often above and below fixation. In the two other groups the cue was more likely (80%) to appear in one of the locations. Target location was counterbalanced and unrelated to the cue location. Participants’ task was to report the direction the arrow was pointing in. **(B)** Reaction time results for Experiment 1. Only trials on which the correct answer was given were used for the analysis. On the left we show results for participants that expected the cue either above or below fixation (N=78), meaning that it was sometimes presented in the expected location and sometimes in the unexpected location. Participants were faster on valid than on invalid trials, regardless of their spatial expectations about the cue. For reference, we present the results for participants in the neutral group (N=40) on the right. Error bars represent SEM.

#### Experiment 1

In Experiment 1 we manipulated the likelihood that the cue would appear either above or below fixation. In two groups of participants the cue was most likely to appear respectively above or below fixation. Consequently, participants in these two groups (N=78) encountered both trials where the cue was in the expected location (80%) as well as trials where it was in the unexpected location (20%). In a third group of participants (N=40) both cue locations were equally likely, resulting in these participants experiencing only neutral trials.

The stimulus-onset asynchrony (SOA) between the cue and target was set to 117 ms. The target remained onscreen until a response was given or until 1000 ms after target onset had passed. Trials were separated by a variable inter-trial interval of 750-1500 ms. Participants responded to the arrows by pressing a button on a button box with their left or right index fingers, respectively for leftward and rightward pointing arrows.

In total the experiment lasted approximately one hour. Before starting with the main experimental task, participants received on-screen instructions and performed one practice block of 80 trials. During the practice block participants received on-screen feedback (response correct or incorrect) on a trial-by-trials basis. Subsequently, participants performed 800 trials of the main task divided over 10 blocks. After each block, participants received feedback about their task performance (overall percentage correct and number of late responses) and subsequently there was a short 20s break.

#### Experiment 2

Instead of manipulating spatial expectations about the cue, in Experiment 2 we manipulated temporal expectations. To remove standard temporal links between cues and targets, the task consisted of continuous blocks (duration 5 min) during which cues and targets were presented (Figure 2A). Temporal expectations were manipulated by varying the regularity of cue presentation onset between blocks. In the regular (expected condition) blocks, the cue would be presented every second (1 Hz presentation rate). In the irregular blocks (unexpected condition) cues were presented quasi-randomly every 0.5 – 1.5 seconds, with a uniform distribution over all possible intervals between two consecutive cues. This results in the participant having less (precise) information about the cue onset and the cue-target SOA. Targets lasted 200 ms and were presented quasi-randomly every 1.0 to 2.5 seconds. Importantly, the onsets of cues and targets were determined completely independently of each other. Because the SOA between two targets was longer than between cues, not every cue was followed by a target. Participants used the ‘z’ and ‘/’ (slash) buttons on the keyboard to respond to leftward and rightward pointing arrow targets respectively.

**Figure 2.**
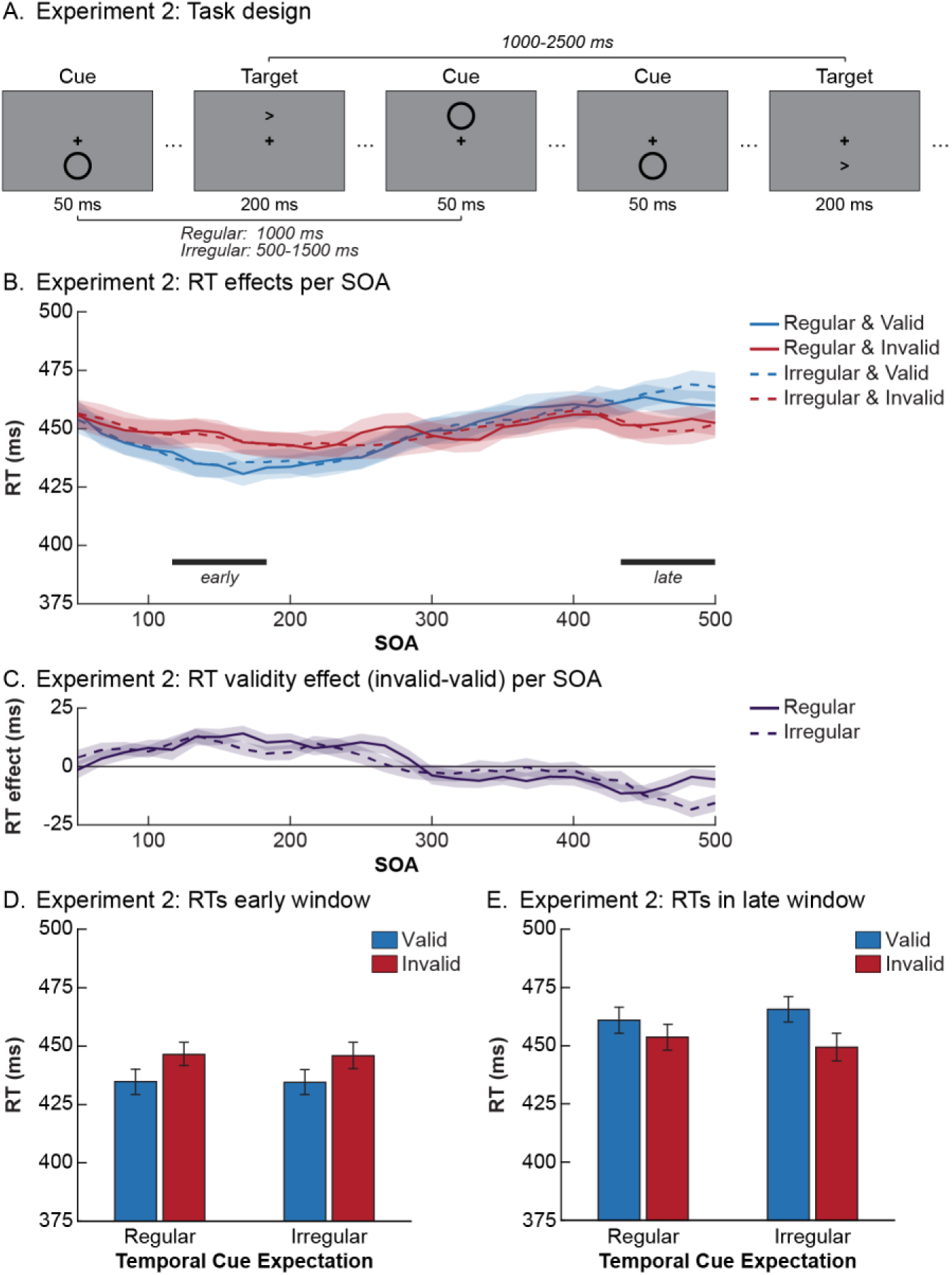
Task design and behavioral results of Experiment 2. **(A)** Example stimulus sequence for Experiment 2. The task consisted of continuous blocks (duration 300s) during which cues and targets were presented. In some blocks cue presentation was regular and hence expectations about cue onset were strong. In other blocks cue presentation was less expectable. Again, participants’ task was to report the direction the target arrow was pointing in. **(B)** The reaction time results and validity effect (invalid-valid) (temporally smoothed) for SOA bins between 50 ms and 500 ms are shown for each of the relevant conditions. Below the graph, the time-periods that were isolated as windows (around 150 ms and 467 ms) for further analyses are marked. **(C)** The validity effect (invalid-valid) results (temporally smoothed) after regularly and irregularly presented cues for SOA bins between 50 ms and 500 ms. The average reaction times per condition from the windows in **(B)** are presented in panel **(D)** for the early window and panel **(E)** for the late window. In both windows there was a significant validity effect that was not affected by temporal cue expectations. Error bars represent SEM.

Participants completed 8 blocks of the task, switching between regular and irregular conditions after every two blocks. The initial condition was counterbalanced over participants. After every block there was a 20s break. If a condition switch occurred, this was explicitly mentioned on-screen at the end of the break. Participants did not receive any feedback during the task. Together with the instructions and the practice block the experiment lasted approximately one hour.

#### Experiment 3

The design of Experiment 3 was largely similar to that of Experiment 2. The most notable difference is that now for each participant the cue location was kept constant throughout the task (Figure 3A). This allowed us to investigate effects of temporal expectations in a context where the distractor is spatially fully predictable. The cue location was counterbalanced across participants. To limit possible carry-over effects, we only switched between the regular and irregular conditions halfway through the experiment. In addition, a short practice block was included at the start of each of the conditions to get participants used to the change in task structure.

**Figure 3.**
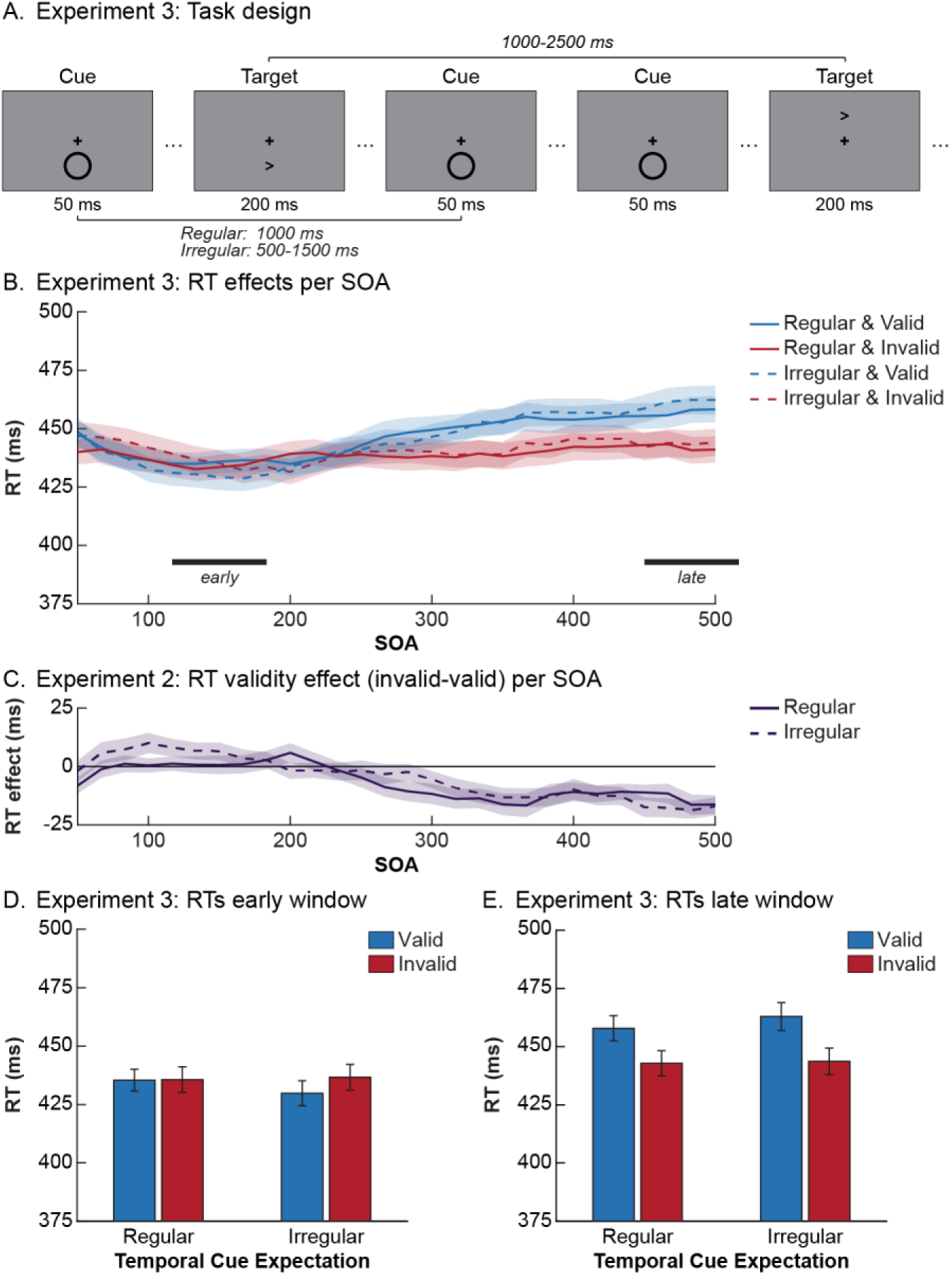
Task design and behavioral results of Experiment 3. **(A)** Example stimulus sequence for Experiment 3. The task that was used was highly similar to that in Experiment 2. The main difference was that for each participant the cue was now consistently presented in one location, making it completely spatially expected. **(B)** The reaction time results (temporally smoothed) for SOA bins between 50 and 500 ms are shown for each of the relevant conditions. Below the graph, the time-periods that were isolated as windows (around 150 ms and 483 ms) for further analyses are marked. In the early window **(C)** there was no significant bottom-up attentional capture, regardless of temporal expectations. The late window **(D)** did show a significant effect, but this inhibition of return effect was not modulated by temporal cue expectations. Error bars represent SEM.

### Behavioral analyses

For all experiments, trials where the participants’ reaction time exceeded 3 standard deviations from the participants’ mean or was below 200 ms were discarded. Furthermore, trials on which no (relevant) response was given within 1 s from target onset were excluded from analyses. The remainder of the trials (Experiment 1: 98.3%; Experiment 2: 96.0%; Experiment 3: 97.0%) was exported to JASP [32] in order to perform statistical analyses. For all experiments, we analyzed the data using a combination of both frequentist statistics and their Bayesian equivalents [33,34].

#### Experiment 1

For the statistical analyses of Experiment 1 we performed a 2 × 2 repeated-measures ANOVA with the factors Validity (valid, invalid) and Expectation (expected, unexpected) for the reaction times. This analysis only considered participants that had expectations about the most likely cue location (N=78). Participants in the neutral condition were a priori included only as a reference group, to be able to interpret possible performance differences as either gains or losses in performance with respect to an expectation-neutral context. In addition to frequentist analyses, we also computed Bayes Factors for all relevant comparisons. As we were specifically interested in the interaction between expectations and the bottom-up validity effect, the Bayes Factor (BF) for a model with only the main effects was compared to the BF for a model with the main effects and the interaction. The ratio of the BF values then quantifies the evidence for including the interaction term in the model, and hence can be interpreted as evidence for or against the existence of an interaction between the two experimental factors. BF ratios will converge either to infinity when a model including the interaction explains the data better, or to zero when it explains the data worse than a model with only main effects. If the ratio is close to one, this indicates that both models are equally likely and that there is not enough evidence for either conclusion. We use the conventions from Jeffreys [35] to interpret the evidence in our Bayesian analyses.

#### Experiment 2 and 3

For each target presentation, we identified the temporal distance between the target and the closest preceding cue stimulus. This generated a total of 90 bins, spanning a cue-target SOA between 0 and 1500 ms, in 17 ms steps (SOAs above 1000 ms were infrequent, <10% of trials). If no response was made within a 1000 ms interval following the target, the trial was classified as a “miss” trial.

Because the number of trials per was low (due to the large amount of possible SOA bins; on average 20.6 trials per SOA bin in Experiment 2 and 21.0 trials per SOA bin in Experiment 3), we smoothed the data in the temporal domain by applying a sliding window over all SOA bins of interest (SOAs below 500 ms). For every relevant combination of conditions, the data we ascribe to an SOA are computed as a weighted average from that SOA and the two SOA’s preceding and following it (0.1*SOA_-2_+0.2*SOA_1_+0.4*SOA_0_+0.2*SOA_+1_+0.1*SOA_+2_). Trials with an SOA between 0 ms and 50 ms were excluded from the analyses because on those trials cue and target presentation overlapped in time.

Based on the smoothed data, we identified two SOAs of interest: (1) an early maximally facilitatory validity effect; and (2) a late maximally inhibitory validity effect. We then averaged the (non-smoothed) data from the selected SOA and two SOA’s preceding or following it to create an early and a late window of interest. Subsequently, we tested for an interaction between expectation and the bottom-up validity effect by performing a 2 × 2 repeated-measures ANOVA with the factors Validity (valid, invalid) and Expectation (expected, unexpected) in each of these windows. In addition, as in Experiment 1 the Bayesian equivalent of the ANOVA was performed.

## Results

### Experiment 1: Do spatial expectations affect bottom-up attention?

In the first experiment we examined whether expectations about the spatial location of a cue modulate the ensuing attentional capture by this cue. In an exogenous cueing task [30], cues and targets were presented above or below fixation. This resulted in valid trials (cue and target in same location) and invalid trials (cue and target in opposite locations). We manipulated the likelihood that the cue would appear at either of the two possible locations between three groups of participants. One group was a neutral control group in which both cue locations were equally likely. In the other two groups the cue was most likely (80% of trials) to appear either above or below fixation. Consequently, those participants encountered both trials where the cue was in the expected location as well as trials where it was not.

In Figure 1B we plot the reaction time results for Experiment 1 for trials on which participants gave the correct answer. Higher reaction times on invalid than valid trials indicate there was attentional capture by the cue (RT difference=29.31 ms, F_1,77_=216.53, p<0.001, η^2^=0.738). Importantly, however, this validity effect was not modulated by the spatial expectation participants had about the cue (F_1,77_=0.753, p=0.388). The evidence against the existence of this modulation is moderate (BF_01_=4.40). While spatial expectations did not modulate attentional capture, there was an overall reaction time benefit for trials where the cue was in the expected location compared to the unexpected location (RT difference=8.87 ms, F_1,77_=84.56, p<0.001, η^2^=0.523). Accuracy was close to ceiling level (98.21±1.21% correct) and hence we did not test for effects on percentage correct.

Together, these results suggest that our manipulation of spatial expectations modulated overall behavioral speed and accuracy but did not result in a modulation of bottom-up attention capture. While this suggests that expectations do not interact with bottom-up attention, there are possible alternative explanations. Most notably, because targets and cues are often presented in the same location, and with only a short (117 ms) and predictable time delay between the two, suppressing processing at the cue location would possibly hamper target processing. Moreover, since the cue was temporally predictive of the target (fixed SOA), its presentation was informative about target onset and hence attending the cue at all times may have been useful for target perception. These considerations inspired Experiment 2, in which we examined whether cues still elicit bottom-up attention when they are temporally predictable but no longer predictive of when targets occur.

### Experiment 2: The effect of temporal expectations on bottom-up attention

Like in the previous experiment, we were interested in the effect of expectations on bottom-up attention. However, contrary to Experiment 1, in Experiment 2 we examined the influence that temporal expectations about cue onset have on subsequent attentional capture by this cue. Moreover, in contrast to Experiment 1 (and almost all experiments on bottom-up attention) in Experiment 2 the cues did not predict when target stimuli occurred. In an adjusted paradigm with longer blocks of visual stimulation (see Methods and Figure 2A), the onset times of cues and targets were manipulated independently. Consequently, cue and target presentation were now unrelated in both the temporal and spatial domain. Temporal expectations about the cue were manipulated by varying the regularity of cue onset between different blocks. In half of the blocks the cue was presented rhythmically (every 1000 ms), supposedly resulting in strong temporal expectations about cue onset. In the other blocks temporal expectations were weaker because the timing of the cue was more variable (500-1500 ms).

Because the number of observations per SOA was low, we temporally smoothed the data over the different SOA bins (Figure 2B). Subsequently, we identified the time-points where numerically the overall validity effect (RT difference) was maximally positive or negative. By averaging data from the five SOAs around those time-points we created an early and late window where bottom-up attention affects were most prominent. The windows can be interpreted as resulting from initial bottom-up attentional capture (early window), followed by inhibition of return (late window; [36]). In these windows, we tested for modulations by temporal expectations (Figure 2C and 2D).

Like in Experiment 1, we observed strong evidence for bottom-up attentional capture as indexed by the validity effect (early window: RT difference=10.54 ms, F_1,60_=24.77, p<0.001, η^2^=0.292; late window: RT difference=-11.18 ms, F_1,59_=15.44, p<0.001, η^2^=0.207). Note that this effect is to be anticipated, since we chose our windows based on the validity effect size. Again, temporal expectations did not modulate bottom-up attentional capture, as shown by the absence of an influence of cue onset regularity on the validity effect (early window: F_1,60_=0.02, p=0.883; late window: F_1,59_=2.85, p=0.097). The evidence against such modulations was moderate for the early window (BF_01_ = 5.14). For the late window, likely related to inhibition of return, there was only anecdotal evidence (BF_01_ = 2.27), suggesting the study’s power was not sufficient to make any strong claims about effects in this time window. Unlike Experiment 1, there was no significant main effect of expectation on reaction time (early window: F_1,60_=0.45, p=0.504; late window: F_1,59_=0.07, p=0.793). Because overall participants’ task performance was close to ceiling (94.00±3.30%), thus we did not test for differences in accuracy between conditions.

### Experiment 3: The role of temporal expectations in bottom-up attention for spatially predictable stimuli

In Experiment 2, the cues in the regular condition were temporally predictable but spatially unpredictable, i.e. they could equally likely occur above or below fixation. This may have reduced the cue predictability.

In this final experiment, we created a condition in which cues were both spatially and temporally fully predictable. Within each participant the cue now always appeared in a single location and thus was always spatially expected. After defining the windows of interest (similar analysis pipeline as Experiment 2; Figure 3B), we again tested whether temporal predictability modulated the bottom-up attention effects (Figure 3C and 3D). As in the previous experiments, no such modulations were found (early window: F_1,55_=1.40, p=0.241; late window: F_1,57_=0.96, p=0.331). The evidence against these interactions was moderate for both windows (early window: BF_01_=3.07; late window BF_01_=3.91). Similar to Experiment 2, there were no main effects of temporal cue expectations on reaction times (early window: F_1,55_=0.09, p=0.763; late window: F_1,57_=1.21, p=0.276). Surprisingly, while there was significant inhibition of return (late window validity effect: RT difference=-17.11ms, F_1,57_=30.38, p<0.001), there was no overall attentional capture in this experiment (early window validity effect: F_1,55_=0.09, p=0.763) and the evidence against the existence of such an effect was moderately strong (BF_01_=3.37). As in the previous experiments, participants’ overall task performance was near-ceiling (95.50±2.72%).

### Comparisons between experiments

Because the experiments differed markedly in the expectations participants had about cues and the context in which those cues were presented, we compared the validity effect sizes between experiments. Therefore, we performed two post-hoc analyses in which we directly compared reaction time validity effects between experiments. To ensure maximal comparability, for Experiment 2 and 3 we take the validity effect at an SOA of 117 ms (after smoothing). First, we compared Experiment 1 (only neutral trials; Figure 4A) and Experiment 2 (Figure 4B). In terms of spatial expectations both are comparable (cue 50% in each location), but the experiments differ strongly in the temporal context in which stimuli are presented. Most notably, in Experiment 1 the SOA was fixed at 117 ms, while in Experiment 2 it was variable and unpredictable. Therefore, in Experiment 1 the cue was a good temporal predictor of target onset. A comparison of the validity effects by means of an independent samples t-test shows a significantly smaller validity effect in Experiment 2 than in Experiment 1 (t_101_=4.47, p<.001, d=.904). This is possibly explained by that fact that in Experiment 2 there was (1) less information about the onset of the target and (2) a lower likelihood (not after every cue) a target would appear at all.

**Figure 4.**
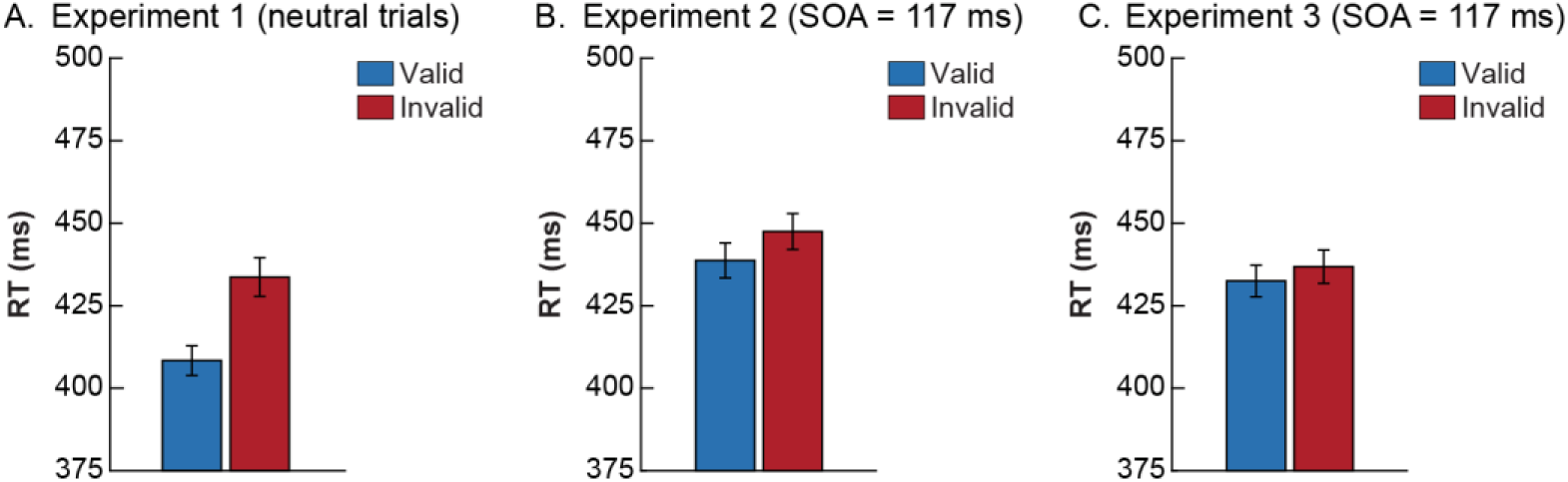
Comparing overall validity effects between experiments. For each of the experiments, we display the average reaction times on valid and invalid trials for the selections of trials that were used to compare validity effects between the experiments. **(A)** Only the neutral condition of Experiment 1 (control group) was used. The SOA in this experiment was fixed at 117 ms. For experiment 2 **(B)** and 3 **(C)** we only used the data we obtained for the SOA of 117 ms after smoothing the SOA time courses. Error bars represent SEM.

Second, the absence of an initial bottom-up capture effect in Experiment 3 (Figure 4C) could be potentially explained by the perfect predictability of cue locations in the experiment. Because in contrast to Experiment 3 cue location was unpredictable in Experiment 2, the comparison between those experiments can be used to test this idea. An independent samples t-test showed that there was no significant difference between the validity effects (t_119_=1.182, p=.240), suggesting that the validity effect was roughly the same in both Experiments. Therefore, we did not find evidence that the bottom-up validity effect is modulated by spatial expectations about the cue. This conceptually replicates our findings of Experiment 1. However, it should be noted that there is only anecdotal evidence (BF_01_=2.75) that the validity effects of Experiment 2 and 3 were of equal size and these results should thus be interpreted with caution.

## Discussion

In a series of three behavioral experiments we investigated whether bottom-up attentional capture is modulated by stimulus expectations. We did not observe empirical support for this idea. On the contrary, in all experiments we found moderately strong evidence that the bottom-up validity effects were of comparable size when cue stimuli were expected, compared to when they were not (or less) expected.

The fact that participants were slightly faster on expected compared to unexpected trials in Experiment 1 suggests that participants did form prior expectations, which had a sizeable influence on behavior. Based on studies showing that stimuli embedded in regular streams are better detected [27], we anticipated to see a similar main effect of expectation for the temporal paradigm in Experiment 2 and 3. Contrary to our predictions, we did not observe this effect, even though regular and irregular blocks were visibly different and every switch between conditions was explicitly marked in the participants’ instructions. Other studies have used similar paradigms with streams of stimulation to investigate effects of temporal expectations and did find effects of temporal regularity [20,37]. Therefore, it appears plausible that temporal expectations about distracting cues do not generally influence processing of target stimuli, when distractors are temporally uncoupled from target stimuli, and target stimuli are salient.

Assuming that our manipulations of expectations actually instantiated priors in our subjects, the absence of any significant interaction between expectations and bottom-up attention in our experiments is surprising, because it contradicts the hypothesis that unexpected or surprising events capture attention (more) [11,13,14]. Interestingly, two recent studies observed that distractor predictability can modulate the amount of attentional capture in a visual search task, where target and distractor are presented concurrently [38,39]. The most notable difference in task design between their study and ours, is that in their paradigm distractors and targets were presented simultaneously on different locations of the screen, resulting in direct attentional competition between the stimuli. This competition can be biased by predictability. In contrast, in our studies we examine the consequences of presenting a salient cue stimulus in isolation on subsequent visual processing at that location. We find that the attention grabbing properties of such a cue are not modulated by predictability. In line with our findings, a recent study by Southwell and colleagues has also suggested that regular and random streams are equally salient [28]. Moreover, our results are in line with the stimulus-driven account of bottom-up attention [2,40], in which it is assumed that the initial capture of attention is automatic and independent from top-down factors. However, it must be noted that even within the stimulus-driven account there would have been room for expectations to suppress the effects of distracting inputs (i.e., the uninformative cues) at later processing stages [4,21].

One conceivable alternative explanation for the absence of effects of expectation on attentional capture is that the tasks we used were too simple. Participant’s performance was close to ceiling (>90%) in all experiments. As a result, ignoring or suppressing the uninformative cues may not have been required in order to perform well. Indeed, recent studies [22,41] showed that attentional capture was only suppressed if task requirements were such that capture would interfere with target processing. Moreover, a study in macaques showed that modulations of V1 attention responses was larger for tasks that were more difficult [12].

Furthermore, a recent study has suggested that only fully spatially predictable distractors may be suppressed at an early processing stage [42]. This hints at the possibility that our manipulation of expectations, especially in Experiment 1, was not potent enough to influence bottom-up capture. It is possible that with a stronger manipulation of expectations (i.e., making cues even more likely in one condition and less likely in the other) we would have observed a modulation of capture. Still, this cannot fully explain the absence of an effect in Experiment 1: even in Experiment 3 when cue stimuli were perfectly predictable in terms of timing, location and visual characteristics, no significant modulation of the validity effect was found.

It is noteworthy that ignoring or suppressing the uninformative cues in our tasks may generally not have been a useful strategy. It is conceivable that participants did not inhibit the cue location at any point in a trial, because a target would often (50%) be presented in the same location with only a short time-delay. As a consequence, ignoring one location systematically would be detrimental to target detection. Moreover, the fixed SOA in Experiment 1 resulted in cue onset being perfectly predictive of target onset time. Hence, paying attention to an informative cue, instead of trying to ignore it, was actually a viable strategy. Consequently, in all experiments the tasks we used may have had factors that made participants attend cues (instead of ignore them). It is possible that the fact that task set did not optimally support suppression of uninformative cues obscured any potential effects in our tasks [9].

We observed a significant difference in the amount of attentional capture between Experiment 1 and Experiment 2. While it is difficult to directly compare both experiments because they differed on several dimensions, a likely candidate explanation for this difference is the fact that in Experiment 2 there was more (temporal) uncertainty about the onset of the targets, as well as an overall lower likelihood of target appearance. It is conceivable that participants deploy a different strategy in Experiment 2 compared to Experiment 1, in which they focus more on the targets and less on the cues (because those are less/not informative), which in turn leads to it having less influence on subsequent target processing. Future studies are required to test this idea.

In conclusion, we did not find evidence for modulations of bottom-up capture by spatial or temporal expectation. We therefore conclude that, at least in the exogenous cueing tasks we used, bottom-up attentional capture was not altered by prior knowledge about the location or time point of the distracting input. This highlights the automatic and involuntary nature of bottom-up attention, and calls into question perceptual surprise as an explanation for bottom-up attention. Future research may use more difficult tasks in which the relationship between targets and distractors can be more carefully controlled. In addition, we believe electrophysiological studies could possibly disentangle the effects of expectation and attention and precisely point at their interactions with high temporal resolution.

## Author contributions

EM, FK, SvG and FdL developed the concept for the set of experiments. All authors contributed to the study design. EM, FK, and LB collected and analyzed the data. All authors were involved in writing the paper and approved the final version of the manuscript.

## Acknowledgements

We thank Joey Zhou, Mariya Manahova and Dirk van Moorselaar for valuable comments on a previous draft of this manuscript.

